# Visualisation of ribosomes in Drosophila axons using Ribo-BiFC

**DOI:** 10.1101/706358

**Authors:** Anand K Singh, Akilu Abdullahi, Matthias Soller, Alexandre David, Saverio Brogna

## Abstract

Rates of protein synthesis and the number of translating ribosomes vary greatly between different cells in various cell states. The distribution of assembled, and potentially translating, ribosomes within cells can be visualised in *Drosophila* by using Bimolecular Fluorescence Complementation (BiFC) to monitor the interaction between tagged pairs of 40S and 60S ribosomal proteins (RPs) that are close neighbours across inter-subunit junctions in the assembled 80S ribosome. Here we describe transgenes that express two novel RP pairs tagged with Venus-based BiFC fragments that considerably increase the sensitivity of this technique that we termed Ribo-BiFC. This improved method should provide a convenient way of monitoring the local distribution of ribosomes in most Drosophila cells and we suggest that could be implemented in other organisms. We visualized 80S ribosomes in larval photoreceptors and in other neurons. Assembled ribosomes are most abundant in the various neuronal cell bodies, but they are also present along the lengths of axons and are concentrated in growth cones of larval and pupal photoreceptors. Surprisingly, there is relatively less puromycin incorporation in the distal portion of axons in the optic stalk, suggesting that some of the ribosomes that have started translation may not be engaged in elongation in axons that are still growing.

## Introduction

Ribosomes are ubiquitous molecular machines that translate gene sequences into the thousands of different proteins that make and operate every organism, so ribosomal components are some of the most abundant and evolutionarily conserved macromolecular constituents of cells. Each ribosome is made up of two complex ribonucleoprotein subunits – 40S and 60S in eukaryotes – and the joining of these into 80S functional ribosomes is tightly regulated. Even when cells are replete with ribosome subunits there are physiological situations (*e.g*. during nutrient deprivation or other cell stresses) when relatively few are assembled into protein-translating ribosomes (Hinnebusch, 2014, 2017).

The joining of ribosomal subunits is a multi-step process, requiring the coordinated activity of several initiation factors, occurring each time that translation of an mRNA is initiated (Hinnebusch, 2017; Jackson *et al.*, 2010). In eukaryotes, the first step is activation of the 40S subunit, which starts with its loading with methionine initiator tRNA (tRNAi^met^). The resulting pre-initiation complex then typically attaches to the 5’ end of an mRNA and scans its 5’UTR until the initiation codon is recognised by base pairing between the anticodon of tRNAi^met^ and an AUG start codon (Kozak, 1989). Once tRNAi^met^ is base-paired with the AUG and is precisely placed in the peptidyl site on the 40S subunit, then the 60S subunit is recruited. The assembled 80S ribosome translocates along the mRNA, catalysing protein synthesis until it reaches a stop codon. It then dissociates and the free subunits become available for new rounds of translation (Dever and Green, 2012).

We have used the Bimolecular Fluorescence Complementation (BiFC) technique to make assembled ribosomes visible in *Drosophila* cells. This is a technique that allows direct detection of diverse types of protein-protein interactions in living cells (Hu *et al.*, 2002; Kerppola, 2008). To do this for ribosomes, one selects a pair of RPs on the surfaces of the individual subunits that only come into close and stable contact when the 80S ribosome assembles, and these RPs are tagged with functionally complementary halves of a fluorescent protein. The two non-functional halves of the fluorescent protein only make a stable contact whilst the 80S ribosome is assembled at initiation, so emission of fluorescence reports that translation initiation has occurred (Al-Jubran *et al.*, 2013).

When we were initially developing the BiFC-based ribosome visualisation technique we tagged several pairs of RPs with either the N-terminal half (YN) or the C-terminal half (YC) of Yellow Fluorescent Protein (YFP). We co-expressed several pairs in *Drosophila* S2 cells and found that only those pairs that come together when the 80S ribosome assembles give rise to ribosomal fluorescence (Al-Jubran *et al.*, 2013). Moreover, the fluorescence was enhanced by translation elongation inhibitors, which stabilise the 80S, and reduced by initiation inhibitors (Al-Jubran *et al.*, 2013). We then designed transgenic flies encoding one such adjacent pair of RPs under UAS regulation (RpS18-YN and RpL11-YC). When these were expressed in salivary glands, a translationally very active tissue that secretes copious amounts of glue proteins (Andrew *et al.*, 2000; Beckendorf and Kafatos, 1976), the tissue showed an intense 80S ribosomal fluorescence signal (Al-Jubran *et al.*, 2013)

We wished to know whether a similar approach could track ribosomes in axons and synapses, and hence serve as a tool for studies of localised translation in the *Drosophila* nervous system (Glock *et al.*, 2017; Holt *et al.*, 2019; Kim and Jung, 2015). Using the available transgenic flies expressing RpS18-YN and RpL11-YC, however, we only detected weak 80S ribosomal fluorescence in the cell bodies of some large neurons, so we sought to improve the sensitivity of this technique we termed Ribo-BiFC. Here we describe an improved version that employs transgenic flies expressing either of two novel RP pairs (RpS18/RpL11 and RpS6/RpL24) that are tagged with BiFC fragments of Venus fluorescent protein (Hudry *et al.*, 2011). These Venus-based reporters greatly improve the sensitivity of the method and reveal clear ribosome signals along the full lengths of axons and at the axon terminals. In larval photoreceptor neurons, which we examined in most detail, intense ribosome signals are also apparent in growth cones. We suggest that these Venus-tagged RP pairs for BiFC, should provide useful research tools with which to monitor the subcellular localisation and trafficking of active ribosomes in most *Drosophila* cells and tissues.

## Results

### BiFC-Venus tagged 80S ribosomes can be detected in axons and growth cones of photoreceptor neurons

The ribosomal protein pairs RpS18/RpL11 and RpS6/RpL24 span inter-subunit potentially contact points, on the surfaces of the ‘head’ and the ‘foot’ respectively, of the 80S ribosome (Figure 1A). We generated UAS-driven *Drosophila* transgenes encoding these proteins that were tagged with complementing fragments of Venus fluorescent protein corresponding to the N-terminal domain (VN, 1-173 aa) and C-terminal domain (VC, 155-238 aa) (Figure 1B). These yield a brighter and more specific BiFC interaction than YFP constructs (Hudry *et al.*, 2011). Moreover, our characterisation in S2 cells indicated that fluorescence from the inter-subunit Venus BiFC complex might be more stable during translation elongation than from the corresponding YFP complex (Al-Jubran *et al.*, 2013).

**Figure 1.**
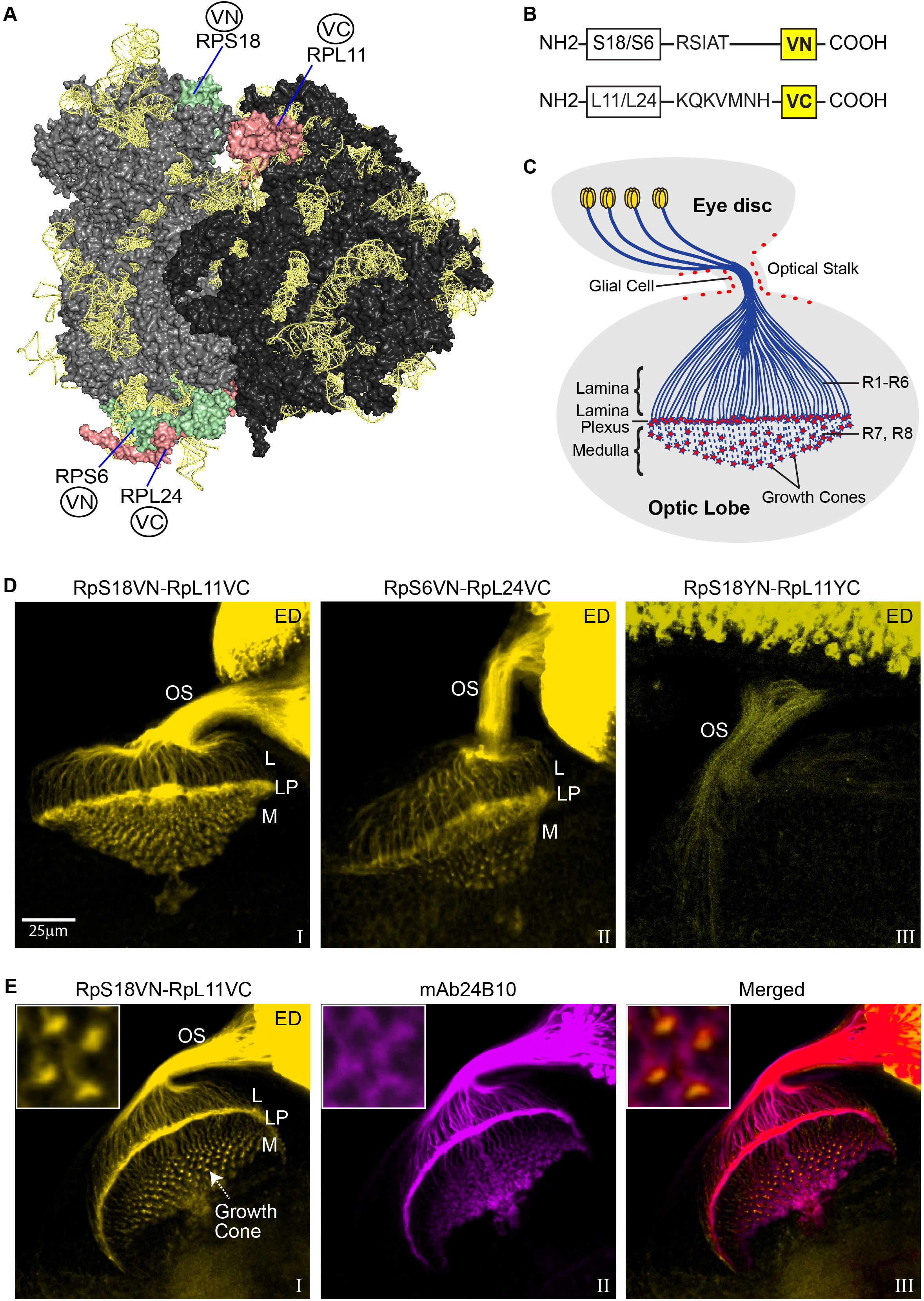
BiFC visualisation of 80S ribosomes in photoreceptors. (**A**) Model of the *Drosophila* 80S ribosome with the two BiFC tagged RP pairs on the small and large subunit highlighted: RpS18/RpL11 and RpS6/RpL24; the image was generated with PyMol using the published high-resolution *Drosophila* 80S structure, PDB file 4V6W (Anger *et al.*, 2013). RpS18 and RpS6 on the 40S are indicated in pale green, RpL11 and RpL24 on the 60S in pale red. (**B**) Diagram of the Bimolecular Fluorescence Complementation (BiFC) constructs with spacer sequences indicated, the Venus VN and VC fragments are shown as yellow boxes. (**C**) Schematic of the eye disc connected by the optic stalk to the brain optic lobe of *Drosophila* larva, showing the photoreceptor cell bodies in the retina (yellow) and their axonal projections into the brain (blue). The photoreceptors R1-R6 project their axons to the lamina region of the brain, while R7 and R8 project their axons further inside to the medulla underneath. The star shapes (red) at the end of axons indicate growth cones. (**D**) Confocal microscopy imaging showing the BiFC signal produced by different transgene combinations expressed in the photoreceptors using GMR-GAL4: RpS18VN/RpL11VC (panel I), RpS6VN/RpL24VC (panel II) and as comparison the YFP-based RpS18YN/RpL11YC (panel III). Labels refer to: OS, Optic Stalk; L, Lamina; LP, Lamina Plexus; M, Medulla. (**E**) Visualisation of the RpS18VN-RpL11VC (yellow, panel I) in tissues in which the photoreceptors are immunostained by mAb24B10 (magenta, panel II), their colocalisation is shown in the merged image (panel III).

We tested the new transgenes in the *Drosophila* larval visual system, which is an excellent model for microscopic visualisation of the axonal projections of neurons. The eye is made up of about 750 ommatidia, each having eight photoreceptor neurons (the R-cells: R1-R8). R1-R6 axons project to a synaptic layer of the brain optic lobe termed the lamina plexus, and R7 and R8 axons pass through the lamina and end in a deeper brain region termed the medulla (Figure 1C) (Mencarelli and Pichaud, 2015). Expression of either of our BiFC-Venus RP pairs in developing eye by using the GMR-GAL4 driver (Freeman, 1996) results in a strong signal. Within the growing photoreceptors, this is brightest in the cell bodies located in the developing eye, but it is apparent along the entire length of the photoreceptor axons, both in R1-R6 (ending in the lamina) and in R7 and R8 (ending in the medulla) (Figure 1D; panel I, RpS18/RpL11; Panel II, RpS6/RpL24). The RpS18/RpL11 pair was used in the experiments described below.

The signal from the Venus-based reporters is much stronger than from the previous YFP-based RpS18/RpL11 transgene pair, which was only apparent in the cell bodies and proximal regions of the axons (Figure 1D, panel III). This was despite the fact that substantial amounts of conventional GFP- or RFP-tagged versions of RpS18 and RpL11, which will report the distributions of free ribosomal subunits as well as assembled ribosomes, are abundantly present throughout the axons (Supplementary Figure 1A).

The neuronal distribution of the signal is confirmed by immunostaining with mAb24B10, which specifically recognizes chaoptin, a GPI-linked cell surface glycoprotein that is present only on photoreceptor neurons and their axons (Figure 1E) (Reinke *et al.*, 1988; Zipursky *et al.*, 1985). There is also intense 80S ribosome signal in enlarged foci at the tips of the R7 and R8 axons in the medulla region (Figure 1E), which is probably in growth cones (Prokop and Meinertzhagen, 2006). Strong signals in photoreceptor growth cones are also apparent during pupal development (Supplementary Figure 2). By comparing the pattern of the 80S signal with that of chaoptin, which mostly stains the periphery of the growth cones (compare insets in Figure 1E), it is clear that the most intense ribosome signal is inside the growth cones. Comparison of the 80S signal with that of mCD8GFP, another plasma membrane marker (Lee and Luo, 1999), which is evenly distributed along the axon (Supplementary Figure1B, panel I vs. panel II), also supports the conclusion that the whole interior of the growth cones must be replete with 80S ribosomes.

We also examined the distribution of 80S ribosomes in other types of neurons by expressing the reporters using different GAL4 drivers (see Material and Methods): D42-GAL4 is expressed in motor neurons (Supplementary Figure 1C, panel I); and DdC-GAL4 and CCAP-GAL4 drive expression in pairs of laterally located neurons that are present in each segment of the brain ventral nerve cord, the axons of which project to the midline (Supplementary Figure 1C, panel II and III, respectively). As in photoreceptor neurons, the fluorescence signals from 80S ribosomes are brighter in the cell bodies, but are apparent along the full lengths of the axons.

### Ribosomes in the distal regions of photoreceptor axons incorporate less puromycin

The classic way to assay for translation is to monitor ribosome-catalysed incorporation of puromycin into the C-terminal of nascent peptides, either radiochemically (Nathans, 1964), or more recently by immunostaining (David *et al.*, 2012; Schmidt *et al.*, 2009). When we incubated salivary glands with puromycin briefly (to minimize diffusion of puromycylated peptides away from translation sites), we saw a good correlation between the 80S BiFC and puromycin signals (Al-Jubran *et al.*, 2013). Puromycin immunostaining has also been recently used to visualise local translation in growth cones of axons that project from Xenopus retinal ganglion cells (Cioni *et al.*, 2019), and mouse brain synaptosomes (Hafner *et al.*, 2019).

We took tissues in which the photoreceptors can be identified by expression either of Venus-based BiFC 80S reporters or of tissue-targetted mCD8-GFP (Lee and Luo, 1999), labelled them and detected puromycylation by immunostaining. Inside the brain the signal was weak and diffuse, and it could not be unambiguously traced to any of the photoreceptor projections or growth cones. However, a clearer pattern was apparent in the eye and optic stalk: it was most intense in the cell bodies in the developing retina and in the proximal regions of their axons (Figure 2A shows distributions in a single longitudinal section of the optic stalk; and Supplementary Figure 3 shows projection images of multiple confocal sections of the same tissue (Panels I-III), and of two other preparations showing similar patterns (Panels IV-IX)). Much of the distribution of the puromycylation signal is similar to that of 80S ribosomes (Figure 2A, panel II and Supplementary Figure 3), but 80S ribosomes are only slightly less abundant in the distal parts of the axons that immunostain weakly for puromycin.

**Figure 2.**
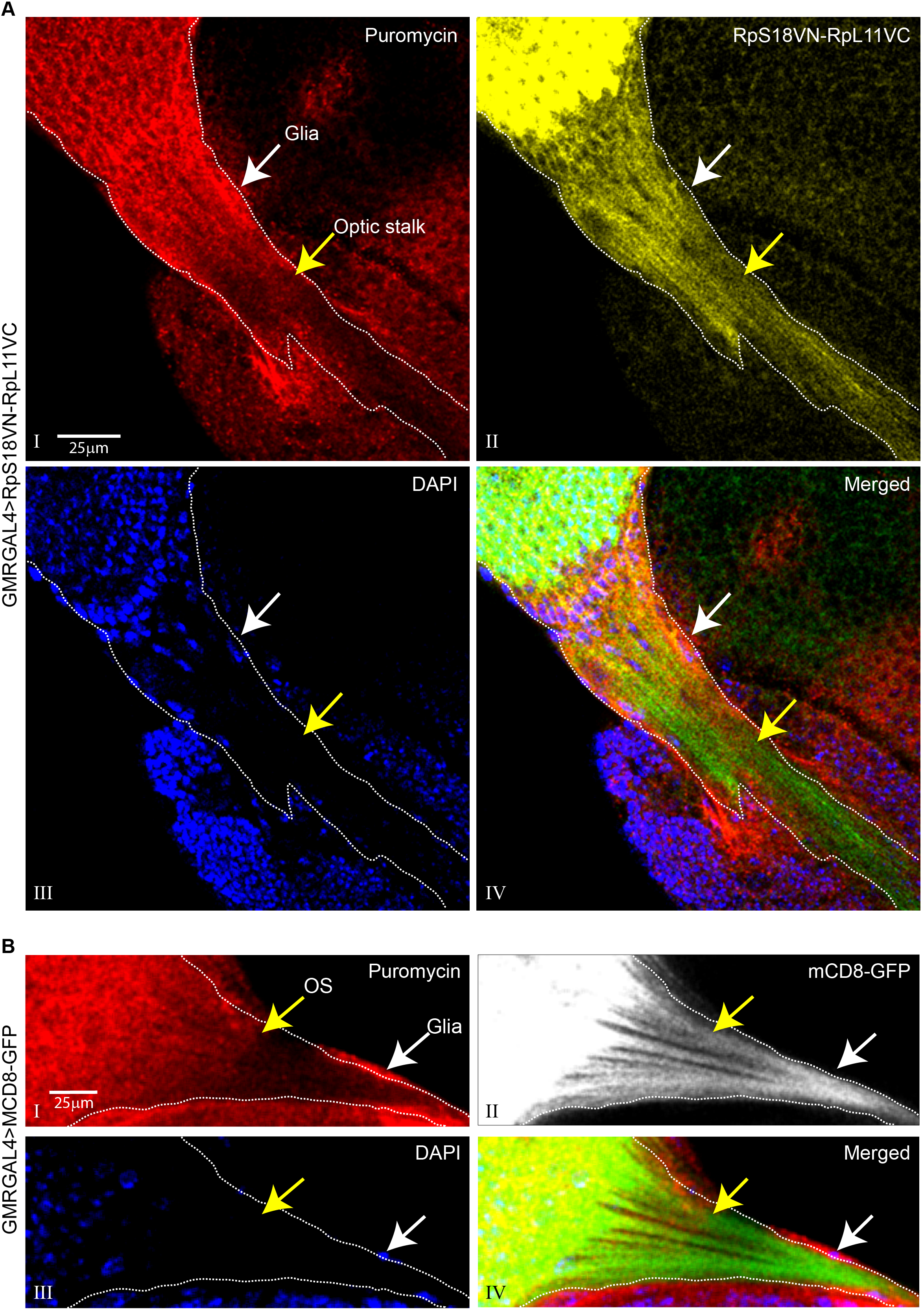
Photoreceptor axons show reduced puromycin incorporation in distal regions. (**A**) Immunocalization of puromycin incorporation (red signal, panel I) in tissues expressing RpS18VN-RpL11VC in the photoreceptors via GMR-GAL4 (yellow, panel II), DAPI staining (blue, panel III) shows the individual nuclei and highlights a monolayer of cells (white arrows), probably glia, surrounding the optic stalk (OS) (yellow arrow); the merged colour image highlights the overlap between the puromycylation and 80S signals in different regions of the photoreceptors (panel IV); the yellow arrow indicates the position of the optic stalk after which there is a reduced puromycylation signal compared to more proximal regions; the BiFC RpS18VN-RpL11VC signal is shown in green instead of yellow in the merged image for better contrast. (**B**) Immunocalization of puromycin incorporation (red, panel I) in tissues expressing GMR-GAL4 driven mCD8-GFP (gray, panel II), DAPI staining shows cell nuclei (blue, panel III); the merged image (panel IV) highlights the relatively more intense green colour in the distal segments of the optic stalk (mCD8-GFP is shown in green for better contrast).

We considered whether the apparent proximal-to-distal gradient of the puromycin signal might be an experimental artifact caused by poor penetration of the antibody into the distal portions of the stalk that extends into the brain. To test this, we examined puromycin incorporation in detergent-permeabilised tissue, in which the photoreceptors were labelled by mCD8-GFP. The puromycin signal was again fainter in the distal regions of the permeabilised axons (Figure 2B). Moreover, there was an intense puromycylation signal in the cells, possibly glia, that surround the entire length of the stalk, indicating that the antibody had free access (Figure 2B, indicated by white arrows). The slower incorporation of puromycin in the distal axonal regions seems therefore not to be caused mainly by a local shortage of ribosomes.

## Discussion

Ribosome activation can be directly visualized by the fluorescence emitted as a result of the interaction between pairs of RPs in different subunits that: a) are tagged with complementary parts of a BiFC-compatible fluorescent protein; and b) are brought into close contact across the junction between subunits when a ribosome assembles. This technique, here named Ribo-BiFC, was previously used to visualise translating ribosomes in *Drosophila* S2 cells and salivary glands (Al-Jubran *et al.*, 2013).

However, our previously described technique was not sensitive enough to visualise ribosomes in all neurons, and here we describe an improved version of the technique. This employs UAS-regulated transgenes that express pairs of neighbouring RPs (RpS18/RpL11 and RpS6/RpL24) that are tagged with BiFC-compatible complementary fragments of Venus fluorescent protein. These new transgenes allow a straightforward and sensitive visualisation of 80S ribosomes in *Drosophila* neurons and clearly detect assembled ribosomes in the axons and growth cones of photoreceptors. We envisage that the sensitivity of this method could be further increased by genetically combining multiple copies of the transgenes we have generated (several P-element inserts are available; see Materials and Methods). These, together with the previously described UAS transgenes encoding individual GFP or RFP-tagged RPs, should provide useful tools that will distinguish between inactive ribosomal subunits and assembled and actively translating ribosomes in *Drosophila* (Rugjee *et al.*, 2013). We propose that our Ribo-BiFC technique should provide a method to visualize changes in the subcellular distribution of ribosomes during different stages of *Drosophila* development and physiological states that will be technically more straightforward than other recently developed methods (Lee *et al.*, 2016). We detected a correlation between the presence of assembled ribosomes and puromycin incorporation, but some of the ribosomes in distal regions of axons seemed not to incorporate puromycin. These may correspond to ribosomes that are either paused on mRNAs after translation initiation or have significantly lower elongation rates. Ribosome pausing has been proposed to be an evolutionarily conserved mechanism to regulate protein synthesis (Darnell *et al.*, 2018); perhaps a similar regulatory mechanism operates on ribosome-loaded mRNAs present in axons of photoreceptor that are still growing and not yet active in the larval stage (Mencarelli and Pichaud, 2015).

## Material and Methods

### Fly Stocks

Generation or the transgenes expressing the YN and YC YFP BiFC fragments or simply GFP tagged ribosomal proteins (RPs) has been previously described (Al-Jubran *et al.*, 2013; Rugjee *et al.*, 2013). The constructs expressing the RPs tagged with either the VN (1–173) and VC (155–238) fragments were similarly generated, cloned in the pUAST vector (Brand and Perrimon 1993), and transgenic flies produced by P element-mediated transformation of standard *yw* strain (Bestgene). The Fkh-Gal4 transgene was used to drive expression in salivary glands (Henderson and Andrew 2000), GMR-GAL4 expresses in the differentiated cells of the developing eye including photoreceptors (Freeman, 1996), D42-GAL4 expresses in motor neurons (Vonhoff *et al.*, 2013), dDC-GAL4 and CCAP-GAL4 express in different groups of neurons in brain ventral cord (Vomel and Wegener, 2008). The UAS-mCD8 GFP transgene encodes a membrane tethered GFP fusion protein used to visualise cell boundaries (Lee and Luo, 1999).

### Puromycylation and Immunostaining

The brain-eye disc tissues of third instar-larvae from mentioned genotypes were dissected in M3 media and incubated with 50 μg/ml puromycin in M3 media for 1 to 10 min. Tissues were briefly washed with M3 media and transferred in 4% formaldehyde for 10 min. Following washing with PBST (0.1% Triton X-100 in 1X PBS) 3 times, tissues were incubated in blocking solution for 1 hr at room temperature followed by mouse anti-puromycin antibody (David *et al.*, 2012)(5B12, 1:500) overnight at 4°C. The mouse anti-chaoptin antibody (mAb24B10, 1:200, DSHB) was used as a neuron specific marker (Zipursky *et al.*, 1985). Tissues were washed with PBST 3 times and incubated with anti-mouse-Cy3 secondary antibody (1:200) for 2 hrs at room temperature. Following washing the tissues were counterstained with 1 μg/mL DAPI (4–6-diamidino-2-phenyl indole, Sigma-Aldrich) and mounted with PromoFluor Antifade Reagent (PromoKine).

### Microscopy

The immunostaining signals in tissues were initially examined under Nikon Eclipse Ti epifluorescence microscope, equipped with ORCA-R2 camera (Hamamatsu Photonics). High resolution images were acquired using a Leica TCS SP2-AOBS confocal microscope. The images were analyzed with either Nikon NIS Elements or Fiji (Schindelin *et al.*, 2012) and figures were prepared using Adobe Illustrator.

## Acknowledgments

We thank Bob Michell and Yun Fan for critically reading the manuscript and discussions. We thank Stephanie Cartwright and Emanuela Zaharieva for their valuable contribution at the start of the project. Thank you also to Suzana Ulian Benitez, Alicia Hidalgo and Yun Fan for providing fly stocks and experimental advice. Thanks to Alessandro Di Maio and the Birmingham Advanced Light Microscopy (BALM) facility; the fly food facility and Shrikant Jondhale for fly stocks maintenance. This project was funded by a Leverhulme Trust (RPG-2014-291) and BBSRC (BB/M022757/1) project grants, and at its start, Wellcome Trust (9340/Z/09/Z) to SB.

---

**Figure S1.**
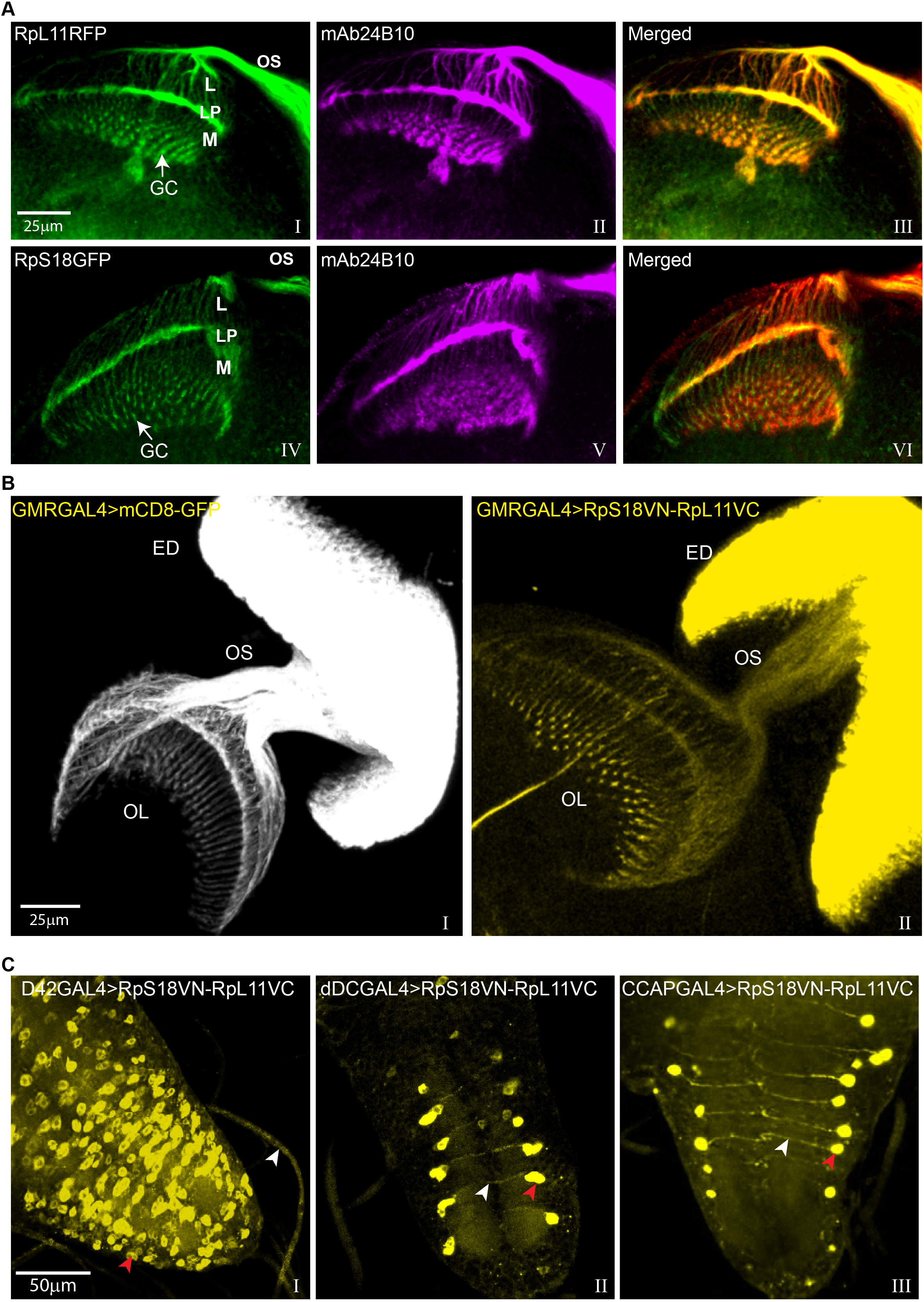
BiFC Visualization of ribosomes in different neurons. (**A**) Localisation of RpL11RFP (green, panel I) or RpS18GFP (green, panel IV) in the R1-R8 photoreceptors immunostained with mAb24B10 (magenta, panel II and panel V); the mAb24B10 is shown in red in the corresponding merged image (panel III and panel VI) for better contrast. Labels refer to: OS, Optic Stalk; L, Lamina; LP, Lamina Plexus; M, Medulla. GC, Growth Cones. (**B**) Localisation of GMR-GAL4 driven mCD8-GFP (gray, panel I) and BiFC signal of RpS18VN/RpL11VC (yellow, panel II) in optic neurons projected from eye disc (ED) via optic stalk (OS) to optic lobe (OL). (**C**) Localization of BiFC signal of RpS18VN/RpL11VC in specific neurons demarcated by the expression of D42-GAL4 (panel I), dDC-GAL4 (panel II) and CCAP-GAL4 (panel III). White arrowheads indicate axons and red ones cell bodies of individual neurons of the ventral nerve cord.

**Figure S2.**
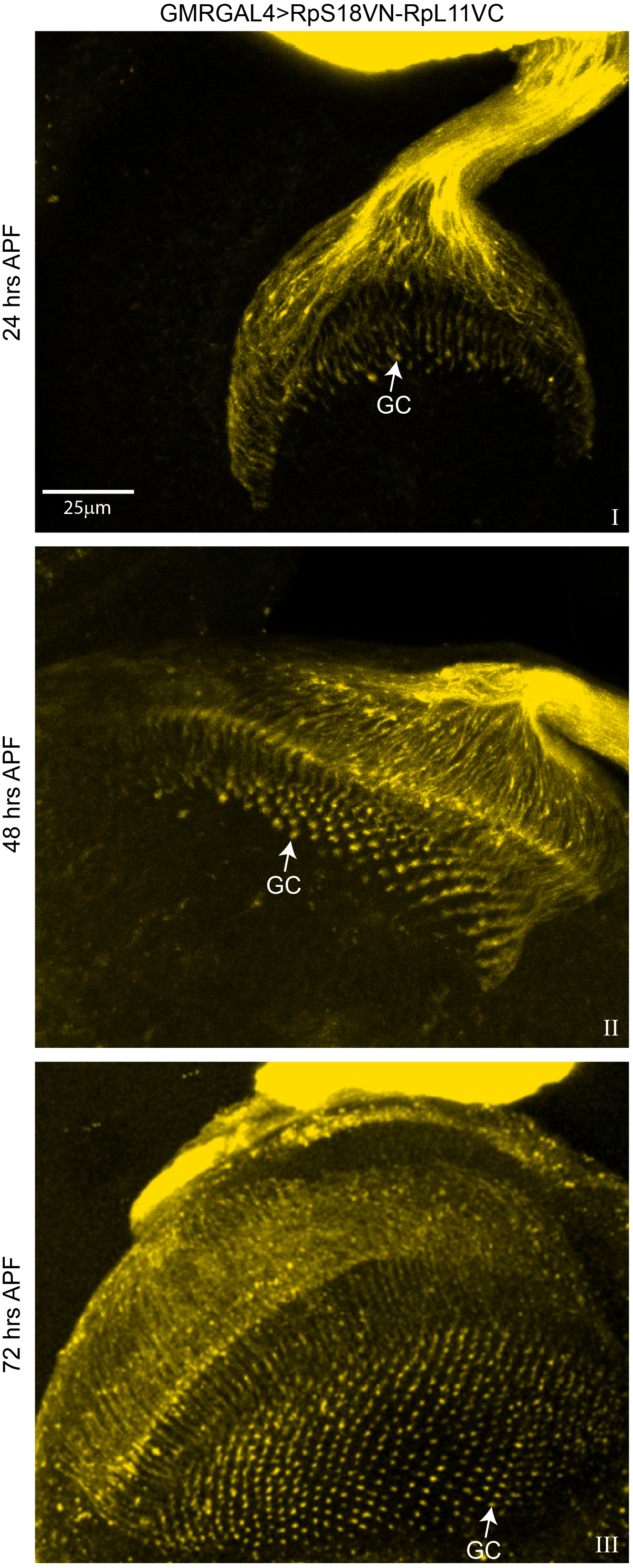
Visualisation or ribosomes in pupal photoreceptor neurons. Localization of BiFC signal of RpS18VN/RpL11VC in photoreceptors at different pupal stages: 24 hrs (panel I), 48 hrs (panel II) and 72 hrs (panel III) after pupa formation (APF). GC refers to Growth Cones.

**Figure S3.**
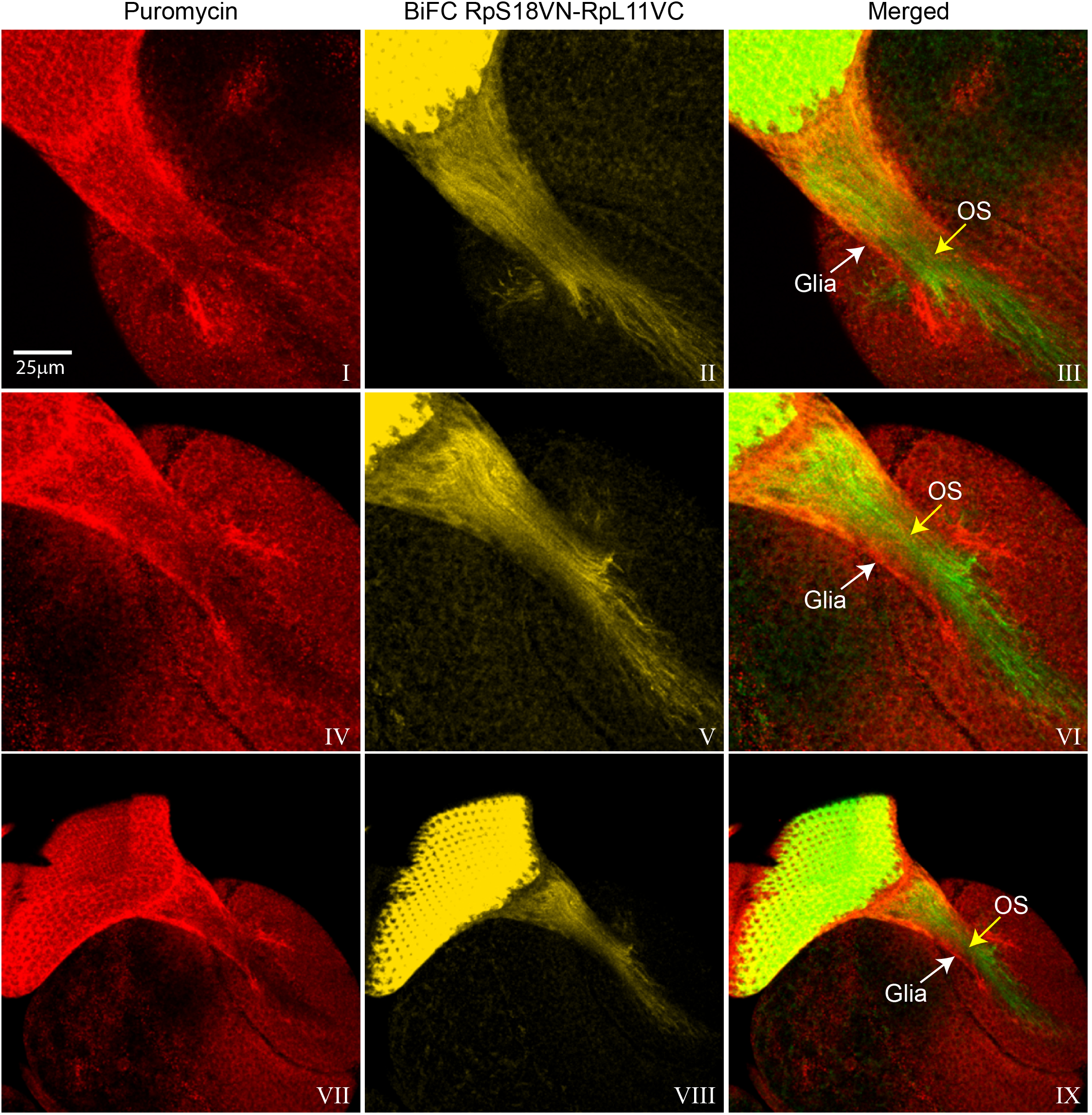
Visualisation of puromycylation sites in the retina and optic stalk. Projection images of the puromycin immunostaining signals (red, panel I, IV, VII) in tissues expressing GMR-GAL4 driven RpS18VN-RpL11VC in the photoreceptors; the BiFC signal (yellow, panel II, V, VIII); the merged image highlights the more intense green colour in the distal segment of the optic stalk (OS) (panel III, VI, IX); the BiFC signal is shown in green for better contrast in the merged image. White arrows indicate a layer of cells, probably glia, surrounding the optic stalks.

